# A NanoLuc luciferase-based assay enabling the real-time analysis of protein secretion and injection by bacterial type III secretion systems

**DOI:** 10.1101/745471

**Authors:** Sibel Westerhausen, Melanie Nowak, Claudia Torres-Vargas, Ursula Bilitewski, Erwin Bohn, Iwan Grin, Samuel Wagner

## Abstract

The elucidation of the molecular mechanisms of secretion through bacterial protein secretion systems is impeded by a lack of assays to quantitatively assess secretion kinetics. Also the analysis of the biological role of these secretion systems as well as the identification of inhibitors targeting these systems would greatly benefit from the availability of a simple, quick and quantitative assay to monitor principle secretion and injection into host cells. Here we present a versatile solution to this need, utilizing the small and very bright NanoLuc luciferase to assess secretion and injection through the type III secretion system encoded by *Salmonella* pathogenicity island 1. The NanoLuc-based secretion assay features a very high signal-to-noise ratio and sensitivity down to the nanoliter scale. The assay enables monitoring of secretion kinetics and is adaptable to a high throughput screening format in 384-well microplates. We further developed NanoLuc and split-NanoLuc-based assays that enable the monitoring of type III secretion-dependent injection of effector proteins into host cells.

**Importance:** The ability to secrete proteins to the bacterial cell surface, to the extracellular environment, or even into target cells is one of the foundations of interbacterial as well as pathogen-host interaction. While great progress has been made in elucidating assembly and structure of secretion systems, our understanding of their secretion mechanism often lags behind, not last because of the challenge to quantitatively assess secretion function. Here, we developed a luciferase-based assay to enable the simple, quick, quantitative, and high throughput-compatible assessment of secretion and injection through virulence-associated type III secretion systems. The assay allows detection of minute amounts of secreted substrate proteins either in the supernatant of the bacterial culture or within eukaryotic host cells. It thus provides an enabling technology to elucidate the mechanisms of secretion and injection of type III secretion systems and is likely adaptable to assay secretion through other bacterial secretion systems.

## Introduction

The ability to secrete proteins to the bacterial cell surface, to the extracellular environment, or even into target cells is one of the foundations of interbacterial as well as pathogen-host interaction. Protein export is particularly challenging for Gram-negative bacteria as two membranes of the bacterial cell envelope have to be passed. So far, nine different protein secretion systems, named type I – IX secretion systems (T1SS – T9SS), have been discovered in Gram-negative bacteria (1, 2). Three of these systems, T3SS, T4SS, and T6SS, serve the direct application of effector proteins into target cells of either prokaryotic or eukaryotic origin (3).

Due to its form and function, the type III secretion machine, as used by many enteric pathogens like *Salmonella*, *Shigella*, *Yersinia*, or enteropathogenic *Escherichia coli*, is called injectisome (4). It is composed of a base that anchors the machine to the inner and outer membranes of the bacterial cell envelope (5), of cytoplasmic components that serve in targeting and receiving of substrates (6, 7), of an inner membrane-localized export apparatus performing substrate unfolding and export (8), and of a needle filament through which secreted substrates reach the host cell (9) (Fig. 1A). Injection itself is mediated by a needle tip complex and by hydrophobic translocators forming pores in the host cell’s target membrane (10). Type III secretion is energized by ATP hydrolysis of the system’s ATPase and by the proton motive force (PMF) across the bacterial inner membrane (11). Secretion of substrates follows a strict hierarchy with early substrates building up the needle filament, intermediate substrates forming the needle tip and translocon pore, and late substrates that serve as effectors inside the target cell.

**Fig 1.**
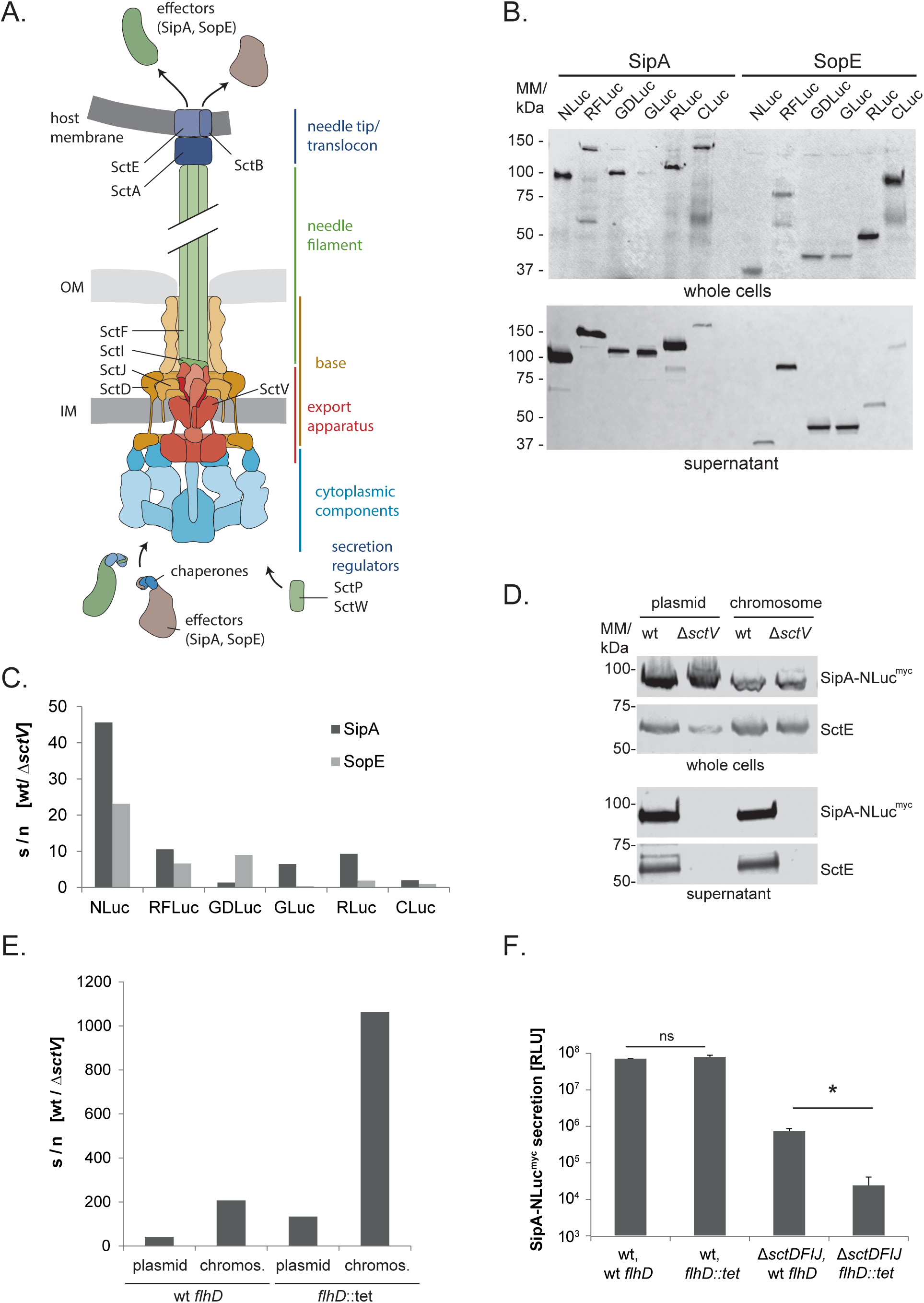
Assessing different luciferases as reporters for type III secretion. (A) Cartoon of the T3SS injectisome. Names or proteins mentioned herein are shown according to the unified nomenclature. The figure is adapted from reference (4). (B) Proteins of whole cell lysates and culture supernatants of *S*. Typhimurium expressing the indicated SipA-Luc and SopE-Luc fusions were analyzed by SDS PAGE, Western blot and Immunodetection with an anti-myc antibody. (C) Signal to noise ratios (wt/Δ*sctV*) of luciferase activities of secreted SipA-Luc and SopE-Luc fusions were graphed. Bar graphs represent the mean S/N of three independent measurements. (D) Immunodetection of SipA-NLuc^myc^ and SctE on Western blot of SDS PAGE-separated culture supernatants and whole cell lysates, either expressing SipA-NLuc^myc^ from a plasmid or from the chromosome. (E) Signal to noise ratios (wt/Δ*sctV*) of luciferase activities of secreted SipA-NLuc either expressed from a plasmid or from the chromosome, each with or without flagella (*flhD*) were graphed. Bar graphs represent the mean S/N of three independent measurements. (F) SipA-NLuc^myc^ secretion in *S*. Typhimurium P_ara_*hilA* and in *S*. Typhimurium Δ*sctDFIJ*, P_ara_*hilA* with and without flagella (*flhD*), respectively. Bar graphs represent mean (± standard deviation) of three technical replicates. Asterisks indicate statistical significance of SipA-NLuc^myc^ secretion assessed by Student’s *t*-test, *: p ≤ 0.05, Abbreviations: Nluc: Nanoluc, RFLuc: Red Firefly luciferase, GDLuc: Gaussia Dura Luciferase, GLuc: Gaussia princeps Luciferase, RLuc: Green Renilla Luciferase, CLuc: Cypridinia Luciferase, S/N: signal to noise, ns: non-significant

While great progress has been made in elucidating assembly and structure of the type III secretion injectisome (12-14), our understanding of its secretion mechanism lags behind, not last because of the challenge to quantitatively assess secretion function. Traditionally, T3SS function is assessed by SDS PAGE, Western blotting, and immunodetection of secreted substrates, either acid precipitated from the bacterial culture supernatant, or analyzed in lysates of eukaryotic target cells (15). This approach is time-consuming, at best semi-quantitative, and lacks sensitivity. To facilitate a simplified analysis of principle secretion, injection, and intracellular localization, several enzyme-linked and fluorescent reporters have been developed (16).

Ampicillin resistance conferred by β-lactamase-fusions secreted into the periplasm was used to monitor the function of flagellar T3SS, which are closely related to T3SS of injectisomes (17). Secretion into the periplasm through partially assembled injectisomes was assessed by using PhoA-fusions, instead (18). While these assays proved very valuable to address some specific questions, monitoring of secretion into the periplasm is only sensible for early substrates as switching to the secretion of later substrates does not occur without an assembled needle. High throughput (HTP) assays for screening of T3SS inhibitors exploited the turn-over of the fluorogenic substrate PED6 by a secreted phospholipase fusion (19), the turn-over of the chromogenic cephalosporine nitrocefin by a secreted β-lactamase fusion (20), and the enzymatic uncaging of the fluorogenic substrate Glu-CyFur by a secreted carboxypeptidase fusion (21, 22).

Likewise, several reporter assays have been developed to assess the injection of T3SS effectors into eukaryotic host cells. Pioneering work by the Cornelis lab exploited the specific increase in intracellular cAMP levels upon injection of effectors fused to a calmodulin-activated adenylate cyclase (Cya) (23). Later, this assay was also adapted to assay injection of effectors by T4SS (24). While the Cya assay showed a very good signal to noise ratio (S/N) of several logs, it was not suitable to monitor injection kinetics or to be adapted for HTP screening because of a tedious cAMP analysis protocol. Widely used to assay injection of effector proteins in T3SS and T4SS is an assay that utilizes the enzymatic cleavage of the FRET-reporter cephalosporin CCF2 by injected β-lactamase-fusions (25). The CCF2 assay facilitated the analysis of injection kinetics and of intracellular accumulation levels of effectors (26). It was also successfully used for HTP high content screening of T3SS inhibitors (27). Real-time observation of injection was achieved by direct fluorescent labeling of tetracysteine motif-tagged effectors (28). However, since this approach requires multidimensional time–lapse microscopy, it is not feasible for routine analysis of effector injection or HTP. Split-GFP technology (29) and self-labelling enzyme tags (30) were successfully used to monitor intracellular localization of effector proteins but both techniques are not optimal for the analysis of translocation kinetics: split-GFP because of a low sensitivity and the slow kinetics of GFP complementation, and the self-labelling enzyme tags because labelling can only be done with effectors that have already been translated before host cell contact.

We aimed to develop a T3SS assay based on effector-luciferase fusions to enable a simple, quantitative, and HTP-compatible assessment of principle secretion and injection. The advantage of luciferase-reporters is a very high S/N and sensitivity. In addition, luciferase-based assays benefit from the lack of product (light) accumulation, simplifying the analysis of secretion and injection rates. We developed a secretion assay utilizing NanoLuc (NLuc) luciferase, an engineered 19 kDa glow-type luciferase from the deep-sea shrimp *Oplophorus gracilirostris* that converts furimazine, emitting blue light (31). The NLuc-based secretion assay allowed quantification of minute amounts of secreted effectors either in the supernatant of the bacterial culture or within eukaryotic host cells. The assay’s ultra-high sensitivity, its wide dynamic range and quick response dynamics qualify it as an enabling technology to elucidate the mechanisms of secretion and injection of T3SS and is likely adaptable to assay secretion through other bacterial secretion systems.

## Results

### Assessment of effector-luciferase fusion proteins as reporters for type III secretion

In order to identify a luciferase compatible with type III secretion through the T3SS encoded by *Salmonella* pathogenicity island 1 (*S*PI-1, T3SS-1), we evaluated six different commercially available luciferases as effector-fused secretion reporters: Cypridinia luciferase (CLuc), Gaussia princeps luciferase (GLuc), Gaussia dura luciferase (GDLuc), NLuc, Renilla luciferase (RLuc), and Red Firefly luciferase (RFLuc) (31-35). We generated translational fusions of the effectors SipA and SopE, respectively, coupled at their C-termini to a luciferase and a myc epitope-tag. The effector-luciferase fusions were expressed from a rhamnose-inducible promoter on a low-copy number plasmid in wild type *S*. Typhimurium and in a secretion deficient mutant (Δ*sctV*). The expression and type III-dependent secretion of the effector luciferase fusions was assessed by SDS PAGE, Western blotting and immunodetection of the myc epitope tag in whole bacterial cells and culture supernatants, respectively, after 5 h of growth. All effector-luciferase fusions could be detected at the expected molecular mass in whole cells and in culture supernatants, indicating their productive expression and secretion (Fig. 1A). CLuc and RFLuc showed additional bands likely corresponding to the cleaved luciferase-myc. In general, SipA-luciferase fusions were secreted more efficiently than SopE fusions. SipA and SopE fusion with CLuc as well as SopE fusions with NLuc and RLuc could only be detected in very low levels in the culture supernatants (Fig.1B).

The activity of the secreted luciferases in filtered culture supernatants of the *S*. Typhimurium wild type and of the Δ*sctV* mutant, respectively, was assessed by luminometry using the specified conditions for each luciferase. The S/N (wild type vs. Δ*sctV*) was highest with effector-NLuc fusions (SipA-NLuc S/N = 45, SopE-NLuc S/N = 22), and, with the exception of GDLuc, always higher for SipA-luciferase fusions (Fig. 1C).

Since the SipA-NLuc fusion showed the best S/N, we introduced SipA-NLuc-myc into the chromosome of a *S*. Typhimurium wild type strain and of a Δ*sctV* mutant for further analysis. First, we compared the expression and secretion of plasmid and chromosome-encoded SipA-NLuc, respectively, and as a reference also of the secreted translocator SctE, by SDS PAGE, Western blotting and immunodetection. SipA-NLuc was expressed well from the chromosome even though, not unexpectedly, at lower levels compared to its expression from the plasmid (Fig. 1D). The extent of T3SS-dependent secretion of plasmid and chromosome-encoded SipA-NLuc was indistinguishable (Fig. 1D).

We next evaluated the S/N of the secreted SipA-NLuc fusion when expressed from plasmid or chromosome by measuring the NLuc activity in filtered culture supernatants of the wild type and the Δ*sctV* mutant. While plasmid-based expression resulted in a S/N = 45, chromosome-based expression even achieved a S/N = 200. The stronger plasmid-based expression may lead to a greater liberation of SipA-NLuc upon occasional cell lysis, compromising the S/N.

Both, injectisomes and flagella possess T3SS for the export of proteins and it has been shown that substrates of one system may be secreted by the other one to a limited degree (36, 37). In order to assess the contribution of the flagellar T3SS to the S/N of SipA-NLuc secretion, we blocked expression of flagella by deleting the gene of the flagellar master regulator FlhD. In the absence of flagella, the S/N of SipA-NLuc secretion increased to 140 when SipA-NLuc was expressed from the plasmid and to 1000 when it was expressed from the chromosome (Fig. 1E). FlhD contributes to the induction of the *S*PI-1-encoded T3SS by indirectly regulating the expression of the the major *S*PI-1 regulator HilA (38), which results in a strongly decreased expression of T3SS-1 and its effectors in the absence of FlhD. To determine whether the improved S/N of SipA-NLuc secretion in the *flhD* mutant resulted from an overall lower expression of the reporter or from preventing secretion through flagella, we also tested SipA-NLuc secretion in a strain expressing chromosome-encoded HilA from an arabinose-inducible promoter (39), thus uncoupling its expression from control by FlhD. In this strain, T3SS-1-dependent SipA-NLuc secretion was identical in the wild type and in the *flhD* mutant (Fig. 1F). However, in the absence of a functional T3SS-1 (Δ*sctDFIJ*), 150-fold lower levels of SipA-NLuc were detectable in the culture supernatant of the strain lacking flagella. These results indicate that about 1% of the SipA-NLuc secretion signal in the wild type strain stems from secretion through the flagellar T3SS (Fig. 1E) and that the increased S/N in the absence of FlhD results from preventing secretion through flagella. Despite the increased S/N in the absence of flagella, we used *flhD* wild type bacteria for most of the work presented herein because of the higher overall signal and because motility appeared to promote growth in a microplate format.

In order to test the versatility of NLuc as secretion reporter, we also constructed fusions with the early T3SS substrate SctP (needle length regulator) and with the intermediate substrate SctA (tip protein). While NLuc compromised secretion and function of SctP when fused to its C-terminus (Fig. S1AB), SctA-NLuc fusions were readily secreted, even when NLuc was placed at different positions within the polypeptide chain of SctA (Fig. S1CD). To overcome the limitation of NLuc in supporting secretion of SctP, we utilized a split-NLuc approach. Split-Nluc is composed of a large fragment (LgBiT, 18 kDa) comprising most of the protein’s beta barrel and of a small fragment with a high affinity to the LgBiT (HiBiT, 1.3 kDa), comprising only one beta strand (40). SctP-HiBiT fusions were successfully secreted into the culture supernatant and strong luminescence was detected when complementing SctP-HiBiT with LgBiT (Fig. S1AB), showing that split-NLuc can serve as a secretion reporter when NLuc fails.

In summary, we could show that luciferases are versatile reporters for T3SS and that effector-NLuc fusions report on secretion with a very high S/N, even in the absence of plasmid-based overexpression.

### Assessment of the sensitivity of the NLuc-based secretion assay

One handicap of the traditional, Western blot-based secretion assay is its low sensitivity that impedes analyzing low culture volumes as required for the analysis of secretion kinetics or for the development of HTP screens.

In order to compare the sensitivity of the Western blot- and the SipA-NLuc-based secretion assays, we made a serial dilution of the filtered supernatant of wild type and Δ*sctV S*. Typhimurium cultures grown for 5 h. In the Western blot-based assay, we could detect the intermediate substrate SctE down to a supernatant volume of 113 µl and the early substrate SctP as well as the late substrate SipA-NLuc down to 225 µl (Fig. 2A). In contrast, using the SipA-NLuc assay, we were able to obtain a stable S/N = 200 down to 195 nl supernatant volume. The S/N even remained above 50 when assaying an equivalent of only 24 nl (Fig. 2B).

**Fig 2.**
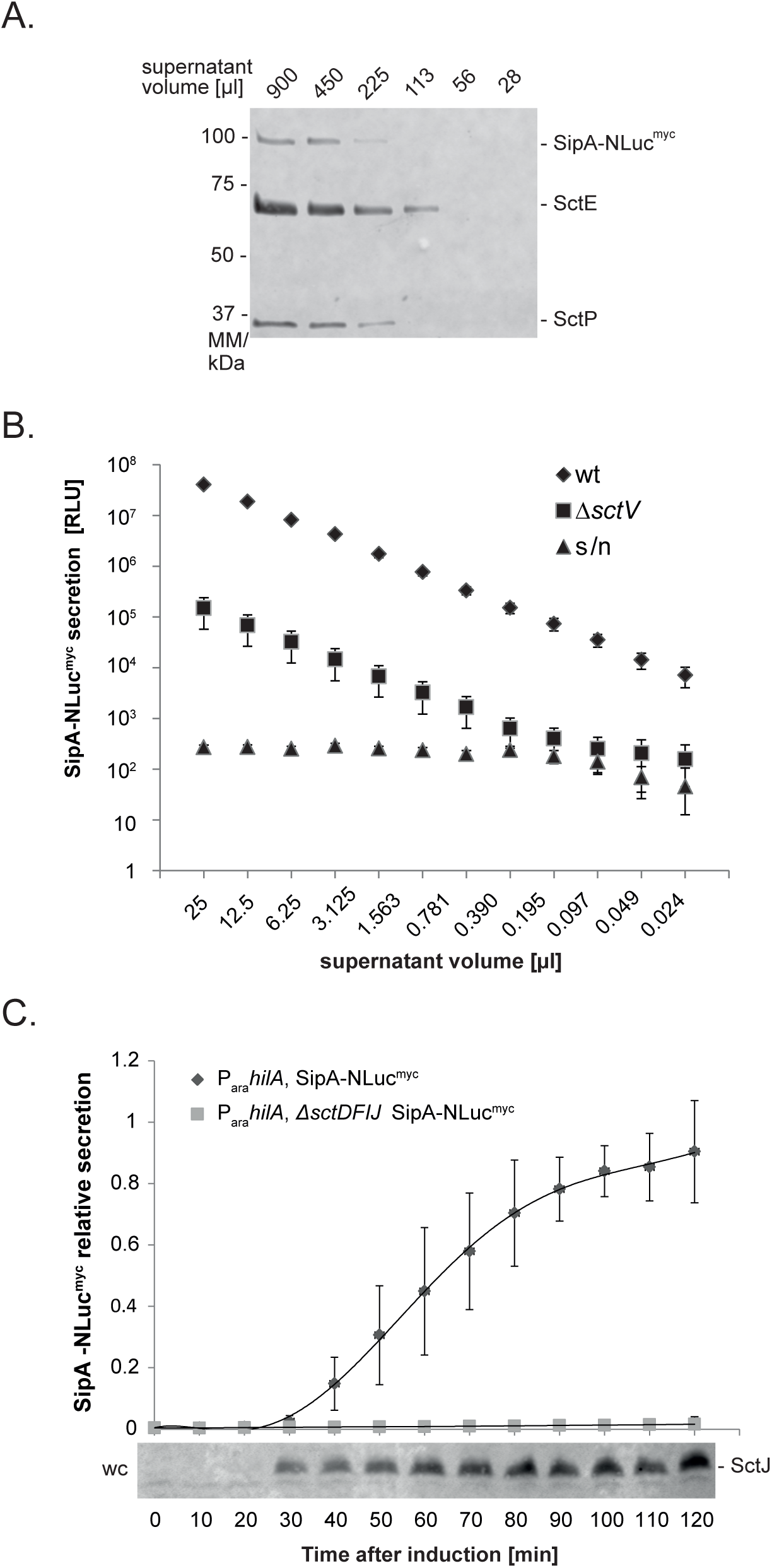
Assessment of the sensitivity of the NLuc secretion reporter. (A) Immunodetection of the T3SS substrates SctP, SctE and SipA-NLuc^myc^ on a Western blot of SDS PAGE-separated, serially diluted culture supernatants. (B) Luminescence of secreted SipA-NLuc^myc^ in serially diluted culture supernatants of the *S*. Typhimurium wild type and a Δ*sctV* mutant. Triangles show the calculated signal to noise ratios for each dilution. Data represent the mean (± standard deviation) of three technical replicates. (C) Normalized luminescence of secreted SipA-NLuc^myc^ at different time points after induction of *hilA* with 0.02% arabinose. Experiments were normalized by setting the maximum luminescence of each experiment to 1. The data points represent mean (± standard deviation) of five independent measurements. At each time point, samples of whole cell lysates were taken for immunodetection of SctJ.

Next, we assessed the performance of the SipA-NLuc assay in monitoring the onset kinetics of type III secretion, which requires very high sensitivity due to the small amounts of secreted material that is initially present. To this end, we grew *S*. Typhimurium harboring arabinose-controlled HilA to an A_600_ = 0.9, after which expression of the pathogenicity island was induced by the addition of 0.02% (w/v) arabinose. Bacterial cells and culture supernatants were collected every 10 min and kept on ice until reading at the end of the experiment. Induction of *S*PI-1 was monitored by Western blot and immunodetection of the base component SctJ in whole cells. It was first observed 30 min after the addition of arabinose (Fig. 2C). Also luminescence of SipA-NLuc was detected in the culture supernatant for the first time 30 min after induction of *S*PI-1 and then luminescence increased steadily to the end of the measurement after 120 min (Fig. 2C). This increase in luminescence correlates directly with SipA-NLuc secretion and is not influenced by NLuc maturation or turn-over as the activity of NLuc remains stable in the culture supernatant over extended periods of time (Fig. S2).

Both experiments, serial dilution and secretion kinetics, prove the superior sensitivity of the NLuc-based over the traditional secretion assay. While the detection of secreted substrate proteins using the traditional assay requires larger volumes and accumulation of substrates in the culture supernatant for an extended period of time, the NLuc assay allows detection of secretion in very small volumes, in brief intervals, and with very short handling times (10 min after collection of supernatant). Our results also show that induction and assembly of the megadalton injectisome is a very quick process that gets bacteria rapidly armed for attack.

### Application example: Harnessing the NLuc secretion assay for high throughput screening

The high sensitivity and the short handling time of the SipA-NLuc-based secretion assay provided an excellent basis to develop a HTP assay for drug screening in a 384-well microplate format.

Centrifugation or filtering is not feasible for separation of bacterial cells and culture supernatant in a microplate format. In order to overcome this problem, we made use of the high-protein binding capacity of the microplates and tested whether secreted substrates would specifically bind to the plate wall after being secreted (Fig. 3A). To this end, 50 µl of *S*. Typhimurium wild type and Δ*sctV* mutant were grown in white high protein binding 384-well plates. Bacteria were washed out of the wells after 5 h of growth using a microplate washer. Then, PBS, NLuc buffer, and NLuc substrate were supplied to each well and the luciferase activity was measured. Using this setup, a S/N = 37 could be achieved (Z’ = 0.8), which is excellent for HTP screening (Fig. 3B).

**Fig 3.**
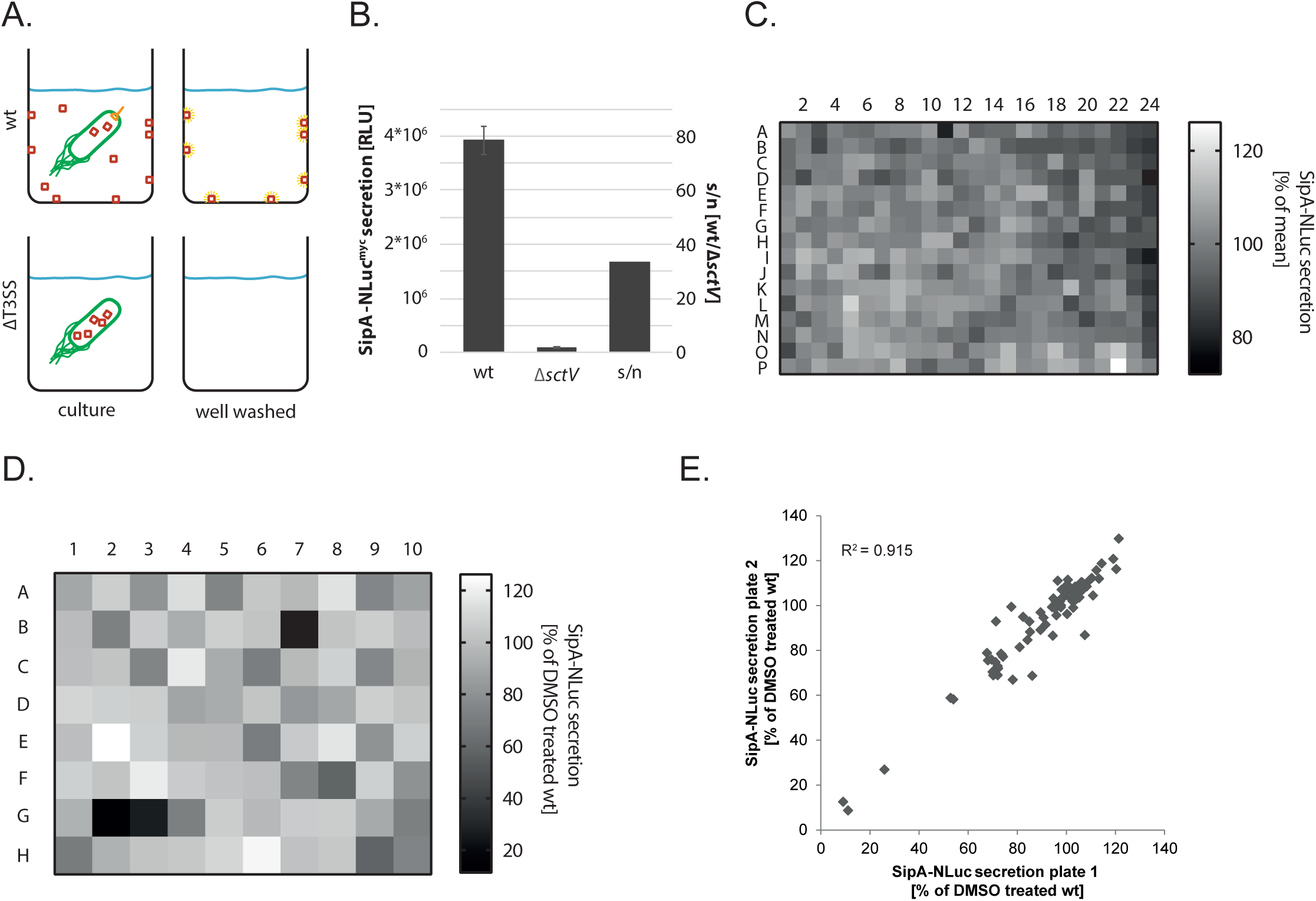
Development of a SipA-NLuc-based HTP secretion. (A) Cartoon of the assay setup. *S*. Typhimurium expressing SipA-NLuc was grown in a 384-well microplate format. Secreted SipA-NLuc bound to the wall of the high protein-binding microplate. Bacteria were washed out and luminescence was measured. (B) Luminescence and signal to noise ratio of secreted SipA-NLuc. The experimental setup was as shown in (A). Bars represent the mean (± standard deviation) of three technical replicates. (C) Signal variation of SipA-NLuc secretion assayed over an entire 384-well microplate as shown in (A). (D) SipA-NLuc secretion in response to treatment with 37 different bioactive compounds, assayed as shown in (A). The DMSO-treated control was set to 100%. The layout of the plate is shown in Table S1. (E) Comparison of the results of two independent compound screens as in (D). The R^2^ value was calculated from a linear regression.

To assess the robustness of this assay and the variation across the plate, we filled an entire 384-well plate with 50 µl of a *S*. Typhimurium, SipA-NLuc culture and allowed it to grow for 5 h at 37°C. Luminescence of secreted, wall-bound SipA-NLuc was assessed after washing out bacteria as described above. The assay proved very robust with a coefficient of variation of 7% over the entire plate and with little edge effects (Fig. 3C, Table S1). We then performed a proof-of-concept inhibitor screen by assessing the effect of a range of 37 different bioactive compounds on the activity of the T3SS in the 384-well format (Table S2, Fig. 3D). Each well of the plate was printed with 0.5 µl of a compound in 100% DMSO, to which 50 µl of a *S*.

Typhimurium, SipA-NLuc culture was added. Again, the culture was allowed to grow for 5 h, after which secretion of SipA-NLuc was assessed by luminometry. The assay showed a highly dynamic response from 10 % to 120 % secretion activity compared to the DMSO-treated wild type control (Fig. 3D). Detection of SipA-NLuc was most strongly reduced by the flavonoids quercetin (30 µg/ml, 90% reduction) and scutellarin (10 µg/ml, 75% reduction), which confirms the previously reported observation that flavonoids target T3SS (22). Also treatment with the 3-hydroxy-3-methylglutaryl (HMG) coenzyme A reductase-blocker simvastatin reduced detection of SipA-NLuc by 44%. Replication of the screen proved a high reproducibility of the assay with a R² of 0.95 (Fig. 3E).

Over all, the SipA-NLuc assay proved to be highly adaptable to a high throughput screening format in 384-well plates, featuring a high S/N, a low error across the plate, a great reproducibility and requiring only short hands-on time.

### Application example: Assessment of the PMF-dependence of type III secretion by the NLuc secretion assay

It has been known for long that secretion through T3SS depends on two sources of energy, on the hydrolysis of ATP by the system’s ATPase (FliI in flagella, SctN in injectisomes) and on the PMF (41-43), which itself is composed of the ΔpH, i.e., the proton concentration gradient across the membrane, and the ΔΨ, the electric potential difference between the periplasm and cytoplasm. The contribution of these two PMF components to T3SS function can be dissected with specific inhibitors. Carbonyl cyanide 3-chlorophenylhydrazone (CCCP) is a PMF uncoupler (ionophore) and discharges both the ΔpH and the ΔΨ by transporting protons through the membrane (44). At acidic pH, potassium benzoate is a weak acid and can enter the membrane and discharge the ΔpH (45). Valinomycin can shuttle potassium ions across the membrane which collapses the electric potential difference ΔΨ (46). Evaluating the contribution of each PMF component to T3SS function requires the careful analysis of secretion kinetics, for which the classical, semi-quantitative Western blot-based secretion assay is not well suited, but for which the NLuc-based secretion assay proved very powerful. To further show this, CCCP, potassium benzoate, and valinomycin, respectively, were added to the bacterial culture at different concentrations, 60 min after induction of *S*PI-1 (for experimental details, please refer to the methods section), while samples of culture supernatants were taken every 10 min for subsequent analysis of the luminescence of secreted SipA-NLuc. While SipA-NLuc secretion progressed over time in the control sample (Fig. 4), addition of the inhibitors lead to sudden changes in secretion kinetics. CCCP blocked secretion instantly, even at concentrations of 5 µM, showing the critical relevance of the PMF for type III secretion (Fig. 4A). Discharching the ΔpH by potassium benzoate resulted in a concentration-dependent instant reduction of secretion (Fig. 4B). At 20 mM potassium benzoate, secretion was completely abolished while it proceeded at 60% of the untreated control in the presence of 5 mM and at 10% in the presence of 10 mM potassium benzoate. Collapsing the electric potential by valinomycin lead to a strongly reduced luciferase signal after 10 min, after which secretion proceeded in a concentration-dependent manner (Fig. 4C): in the presence of 20 µM valinomycin, no significant change in secretion rate was observed, while 40 µM and 60 µM valinomycin, respectively, lead to 70% and 40% secretion of the untreated control.

**Fig 4.**
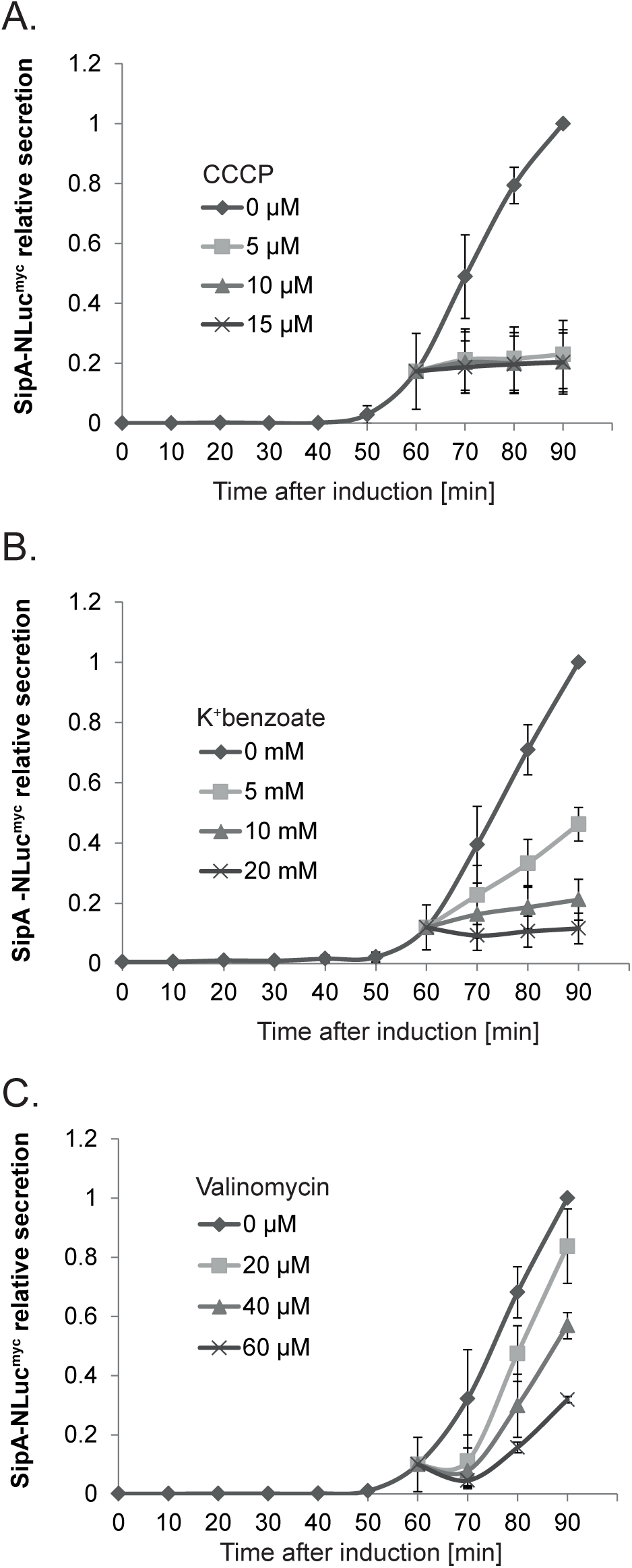
Assessment of the PMF-dependence of type III secretion by the NLuc secretion assay. (A) Normalized secretion of SipA-NLuc in *S*. Typhimurium P_ara_*hilA* after induction of *S*PI-1 by addition of 0.02% arabinose. CCCP was added to a final concentration of 0, 5, 10 and 15 µM, respectively, 60 min after induction of *S*PI-1. (B) As in (A) but addition of K^+^ benzoate to final concentration of 0, 5, 10 and 20 mM, respectively. (C) As in (A) but addition of Valinomycin to a final concentration of 0, 20, 40 and 60 µM, respectively. All data represent means (± standard deviation) of three independent measurements.

These results show that both components of the PMF, ΔpH and ΔΨ, contribute to energizing secretion in the *S*PI-1-encoded T3SS of *S*. Typhimurium. As the PMF-compromising compounds took effect so quickly after treatment, it is highly unlikely that the PMF-dependence of type III secretion is the consequence of a secondary effect of PMF reduction – an issue that could only be resolved with the sensitive and highly time-resolved NLuc secretion assay. These results open the door for further experiments dissecting the role of the different T3SS components in utilizing the PMF.

### Development of NLuc-based host cell injection assays

Assessment of secretion of T3SS substrates into the culture supernatant is very useful for investigating the basic secretion mechanism of T3SS, however the intended biological function of T3SS injectisomes is the injection of effector proteins into host cells. Since the SipA-NLuc-based secretion assay proved to be very sensitive and simple, we attempted to adapt the assay to monitoring the injection of SipA-NLuc into host cells.

In a first and simple approach, we infected HeLa cells in 96-well plates at an MOI = 50 with SipA-NLuc-expressing *S*. Typhimurium, using wild type bacteria and secretion-deficient Δ*sctV* mutants. After infection for 60 min, attached bacteria were gently washed off with PBS using a microplate washer and subsequently, the HeLa cell-associated luminescence was measured using live cell buffer (Fig. 5A). The non-secreting Δ*sctV* mutants (Fig. 5A) showed a HeLa cell-associated luminescence of 8% of the wild type, corresponding to a S/N = 12 (Fig. 5C). To determine whether the HeLa cell-associated signal was truly resulting from injected SipA-NLuc, we assessed injection in a set of mutants that are capable of secreting SipA but incapable of injecting it into host cells: a needle tip-deficient Δ*sctA*, a translocon-deficient Δ*sctEBA*, and a gatekeeper-deficient Δ*sctW* mutant. While secretion of SipA-NLuc into the culture supernatant was increased between 2 and 5-fold in Δ*sctA*, Δ*sctEBA*, and Δ*sctW* mutants (Fig. 5B), which are reportedly unlocked for secretion of late substrates like SipA (47, 48), the HeLa cell-associated luminescence was strongly reduced to 9-24% of the wild type when infecting with these mutants (Fig. 5C). From these results we can conclude that the luminescence signal obtained from infection with wild type *S*. Typhimurium resulted to more than 90% from injected SipA-NLuc and that only little signal may stem from bacteria remaining attached to HeLa cells or to the plate even after washing. Over all, this NLuc-based injection assay proved very useful for the quick and simple assessment of translocation of effectors into host cells by an end-point measurement, however the kinetics of injection cannot be assessed by this assay.

**Fig 5.**
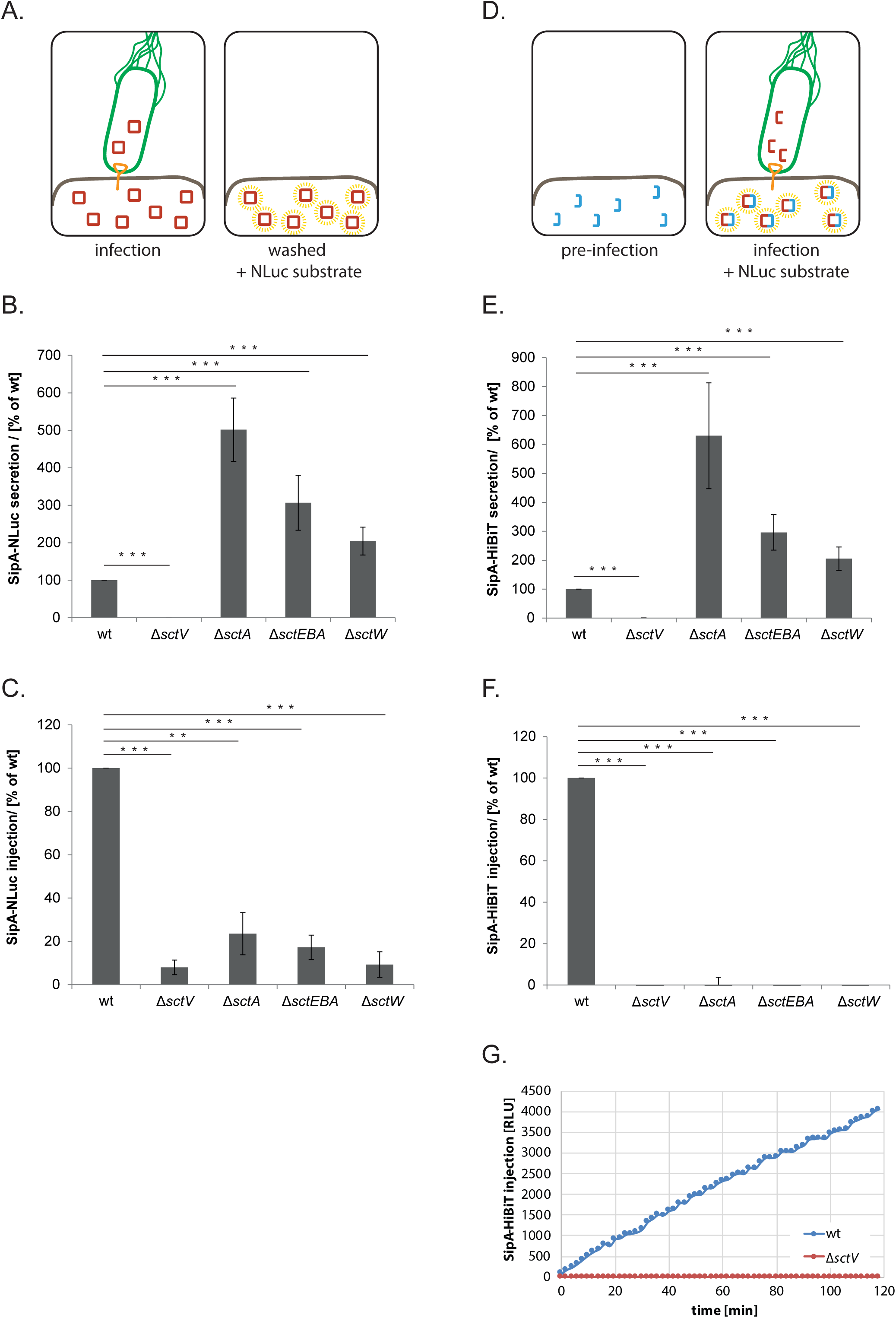
Development of NLuc-based host cell injection assays. (A) Cartoon showing setup of NLuc injection assay. *S*. Typhimurium expressing SipA-NLuc was allowed to infect HeLa cells for 60 min. SipA-NLuc was injected into HeLa cells by use of the T3SS injectisome. Bacteria were washed away using a microplate washer and subsequently NLuc luminescence was measured. (B) Luminescence of SipA-NLuc secreted by the *S*. Typhimurium wild type and indicated mutants in the absence of host cells. The luminescence of the wild type was set to 100%. (C) Luminescence of SipA-NLuc injected into HeLa cells by the *S*. Typhimurium wild type and indicated mutants. The experimental setup was as shown in (A). The luminescence of the wild type was set to 100%. (D) Cartoon showing setup of split-NLuc (HiBiT) injection assay. *S*. Typhimurium expressing SipA-HiBiT was allowed to infect HeLa cells (expressing LgBiT) for 60 min. SipA-HiBiT was injected into HeLa cells by use of the T3SS injectisome. Luminescence of the complemented split-NLuc was measured. (E) Luminescence of LgBiT-complemented SipA-HiBiT secreted by the *S*. Typhimurium wild type and indicated mutants in the absence of host cells. The luminescence of the wild type was set to 100%. (F) Luminescence of SipA-HiBiT injected into LgBiT-expressing HeLa cells by the *S*. Typhimurium wild type and indicated mutants. The experimental setup was as shown in (D). The luminescence of the wild type was set to 100%. (G) Luminescence of SipA-HiBiT injected into LgBiT-expressing HeLa cells by the *S*. Typhimurium wild type and the Δ*sctV* mutant. At timepoint zero, HeLa cells were infected with *S*. Typhimurium after which cells were incubated inside a microplate reader in the presence of NLuc substrate. Luminescence was followed in 2 min intervals. Values of the Δ*sctV* mutant were set to zero for each time point. The results show the mean of technical triplicates. Bar graphs represent mean (± standard deviation) of three independent measurements. Asterisks indicate statistical significance between wt and mutant strains assessed by a Students *t*-test, ***: p ≤ 0.001 **: p ≤ 0.01

To gain a higher specificity for the signal of injected SipA and enable analysis of injection kinetics, we employed the split version of the NLuc luciferase. To this end, SipA was fused to HiBiT while LgBiT was expressed stably by the HeLa cell line. Complementation of LgBiT with HiBiT to a functional luciferase should only occur inside the HeLa cells after translocation of SipA-HiBiT (Fig. 5D). We first tested the secretion of SipA-HiBiT into the culture supernatant by providing LgBiT to the assay buffer. Similar to what was observed for SipA-NLuc, secretion of SipA-HiBiT into the culture supernatant was increased between 2 and 6-fold in Δ*sctA*, Δ*sctEBA*, and Δ*sctW* mutants, respectively (Fig. 5E). However, in contrast to the SipA-NLuc-based injection assay, none of the T3SS mutant strains yielded any detectable luminescence in the split NLuc assay (Fig. 5F), making this assay highly suitable for monitoring the specific injection of T3SS effectors into host cells. This setup even allowed us to follow the kinetics of SipA-HiBiT injection over time directly in a microplate reader (Fig. 5G).

## Discussion

The elucidation of the molecular mechanisms of secretion through T3SS and other bacterial protein secretion systems is impeded by a lack of assays to quantitatively assess secretion kinetics. Also the analysis of the biological role of these secretion systems as well as the identification of inhibitors targeting these systems would greatly benefit from the availability of a simple, quick and quantitative assay to monitor principle secretion and injection into host cells. Here we present a versatile solution to this need, utilizing the small and very bright NLuc luciferase to assess secretion and injection through the T3SS encoded by *S*PI-1 of *S*. Typhimurium. Secretion of a SipA-NLuc fusion showed a very high S/N and sensitivity down to the nanoliter scale, making it exquisitely suited for the assessment of secretion kinetics. In addition, the NLuc-based secretion assay proved highly adaptable to a HTP screening format in 384-well microplates. We further developed NLuc and split-NLuc-based assays that enable the monitoring of T3SS-dependent injection of effector proteins into host cells.

A perfect assay to monitor protein secretion would feature: i) A lack of signal from the unsecreted reporter, resulting in a high S/N. ii) A small reporter that does not interfere with secretion through the secretion system of interest. In case of T3SS, this also includes a not too fast and tight folding inside bacteria as only unfolded protein can be secreted and as the unfolding capacity of the system is not very high. iii) A fast and efficient folding of the reporter outside of the bacterium, guaranteeing fast response dynamics. iv) An intrinsic signal of the reporter, not necessitating an enzyme substrate. v) A high sensitivity. vi) A lack of accumulation of product of the reporter’s reaction. And vii) Be quick, simple, and needing only short hands-on time.

While fluorescent proteins would be desirable secretion reporters as they benefit from an intrinsic signal (and thus do not come with the problem of accumulation of product of the reporter’s reaction), they often suffer from a very slow maturation time and/or insufficient brightness. In addition, fluorescent proteins tend to form very stable β-barrels that block secretion through T3SS (49), excluding them as secretion reporters, at least for T3SS. While the use of split GFP can overcome the limitation associated with tight folding, slow complementation and maturation of GFP compromise its use. The NLuc-based secretion assay as presented herein matches most of the needs listed above. While NLuc lacks an intrinsic signal and requires the addition of a substrate, the analysis of secretion by this assay is not complicated by the overlay of the kinetics of the reporting enzyme and the kinetics of secretion, as it is in other enzyme-linked secretion assays. Instead, the measured signal of the NLuc assay is directly proportional to the amount of accumulated secreted protein. This advantage, together with the superior sensitivity, yield a very high dynamic range of the NLuc secretion assay.

We demonstrated that the NLuc-secretion assay is highly suited to study the kinetics of secretion due to its superior sensitivity. Our simple assay setup only allowed deduction of secretion kinetics from the accumulation of NLuc in the culture supernatant but culturing bacteria in a microfluidics system could enable the direct and on-line reading of secretion into the medium flow through and by this facilitate an even better resolved analysis of the mechanism of secretion.

Our experiments show that secretion of NLuc is supported by fusion to a range of intermediate and late T3SS substrates, even within a polypeptide chain, but fails to be secreted when fused to the early substrate SctP. It is conceivable that the mode of early substrate secretion does not provide a sufficient unfolding capacity to support secretion of NLuc while this seems not a problem when NLuc is fused to intermediate and late substrates. Interestingly, a *Yersinia* SctP-PhoA fusion was secreted (18), pointing either to a higher unfolding capacity of the *Yersinia* T3SS or to a weaker fold of PhoA. We could overcome the limited use of NLuc as secretion reporter for early substrates by using the split-NLuc technology instead. The 11 amino acid-long HiBiT was accommodated well by SctP and it is conceivable that this very small piece allows secretion in most circumstances.

In its current form, the NLuc secretion assay requires the separation of bacteria and supernatant to achieve a good S/N because of the membrane-permeating properties of the NLuc substrate. A membrane impermeant NLuc substrate could overcome this limitation, would make NLuc-based secretion assays even more simple and versatile and increase their robustness due to less steps of handling.

In addition to the points important for a secretion assay, a perfect injection assay would also: i) Feature a high specificity for injected effectors as opposed to secreted but not injected ones. ii) Allow the analysis of injection kinetics. And iii) Allow localization of the injected protein, at best at single molecule resolution.

While fluorescence-based assays proved highly suitable to study the localization dynamics of injected effectors inside host cells, they are very limited in their use to study injection kinetics and are always instrumentation-demanding. The CCF2-based injection assay features simple handling, instead, and proves very useful for the analysis of injection, but suffers from high costs of CCF2 and a low dynamic range. In addition, the product accumulation resulting from the enzymatic activity of the injected β-lactamase complicate the analysis of injection kinetics. The herein-presented NLuc-based injection assays offer very quick and simple analysis, even of injection kinetics, and feature a high dynamic range and sensitivity. While a high-resolution analysis of the localization of the effector-NLuc-fusions inside host cells is not supported by these assays, microscopic setups exist that utilize luminescence for long-duration monitoring of single cells (50), which may become useful for studying the role of individual effectors in bacterial persistence.

As performed herein, cytoplasmic expression of LgBiT will only generate luminescence if the HiBiT of the injected effector also localizes to the cytoplasm. However, the split-NLuc injection assay may also be utilized to analyze the localization and topogenesis of effector proteins inside host cells by targeting LgBiT to specific organelles instead (Fig. 6). Furthermore, complementation of LgBiT by the low-affinity SmBiT instead of the high-affinity HiBiT may provide a useful tool to investigate effector-host protein interactions *in vivo* by bimolecular complementation (51).

**Fig 6.**
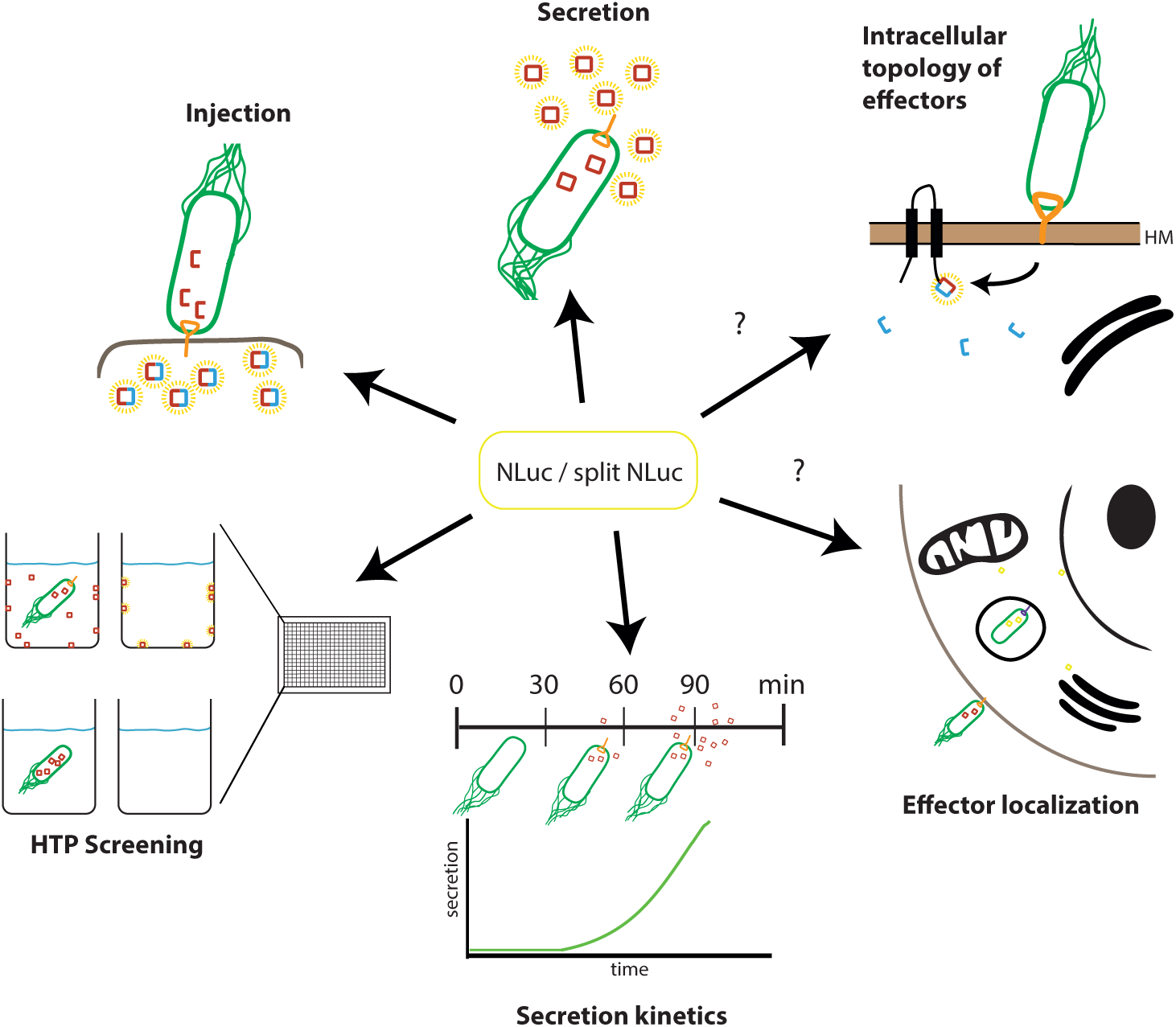
Cartoon summarizing the utilization of the NLuc-based T3SS secretion and injection assays.

In summary, our data show that NLuc-fusions of secreted substrate proteins can be used as a robust, versatile, cheap, simple and quick reporter for T3SS secretion and injection that will enable future in-depth elucidation of T3SS function (Fig. 6). The NLuc reporter is likely to be adaptable to other bacterial secretion systems as well.

## Materials and methods

### Materials

Chemicals were from Sigma-Aldrich unless otherwise specified. SERVAGel™ TG PRiME™ 8–16% precast gels were from Serva. Primers, listed in Table S3, were synthetized by Eurofins and Integrated DNA Technologies. Monoclonal anti-c-myc antibody was from Roche (11-667-149-001). Secondary antibodies goat anti-mouse IgG DyLight 800 conjugate were from Thermo-Fisher (SA5-35571).

### Bacterial strains, plasmids and growth conditions

Bacterial strains and plasmids used in this study are listed in Table S3. All *Salmonella* strains were derived from *Salmonella* enterica serovar Typhimurium strain SL1344 (Hoiseth and Stocker, 1981) and created by allelic exchange as previously described (52). *S*. Typhimurium strains were cultured with low aeration at 37°C in Lennox broth (LB) supplemented with 0.3 M NaCl to induce expression of *S*PI-1. As required, bacterial cultures were supplemented with tetracycline (12.5 µg/ml), streptomycin (50 µg/ml), or kanamycin (25 µg/ml). Plasmids were generated by Gibson cloning (53) using KOD (Novagen) or Q5 polymerase (NEB). Expression of pT10-based plasmids was induced by the addition of 100 µM of rhamnose to the culture medium.

### Western-blot-based secretion assay

Western-blot-based analysis of type III-dependent secretion of proteins into the culture medium was carried out as described previously (39). *S*. Typhimurium was cultured at 37°C for 5 h. For separation of whole cells and cell culture supernatant, the bacterial suspensions were centrifuged at 10,000 × *g* for 2 min at 4°C. Whole cells were directly resuspended in SDS PAGE loading buffer. The supernatant was filtered through a 0.22 µm pore size filter, sodium deoxycholic acid was added to a final concentration of 0.1% (w/v), and proteins were precipitated by addition of 10% trichloroacetic acid (v/v; final concentration) for 30 min at 4°C. After pelleting by centrifugation at 20,000 × *g* for 20 min at 4°C, precipitated proteins were washed with acetone and subsequently resuspended in SDS PAGE loading buffer.

### Luciferase assays

To measure NLuc, RFLuc, Gluc, GDluc, Rluc and Cluc activity of secreted translational fusions, bacteria were grown under *S*PI-1-inducing conditions for 5 h. Culture supernatants were separated from whole bacterial cells by centrifugation for 2 min at 10,000 × *g*. The following buffers were prepared with their substrates according to the manufacturers’ protocols: For Nluc, 25 µl of Nano-glo assay buffer containing furimazine (Nluc working solution, Promega) was added to 25 µl of the culture supernatant. For RFLuc, 30 µl of constituted One-glo assay buffer containing luciferin (Promega) was added to 30 µl of the culture supernatant. For Gluc and GDLuc, 50 µl of the assay buffer containing coelenterazine (Thermo Fisher) was added to 20 µl of culture supernatant. For RLuc, 25 µl of the constituted assay buffer (Promega), in which the substrate was 1:100 diluted, was added to 25 µl of the culture supernatant. For CLuc, a working solution was prepared containing assay buffer and 1:100 of the substrate vargulin (Thermo Fisher). 30 µl of the working solution was added to 10 µl of the supernatant. The luciferase activities were measured in white 384-well plates (MaxiSorp, Nunc), with acquisition settings as recommended by the manufacturers.

### NLuc assay for wall-bound protein

In order to measure wall-bound protein, overnight cultures of *S*. Typhimurium were back-diluted to an A_600_ = 0.1 and 50 µl of the bacterial suspension was transferred to a 384-well microplate (MaxiSorp, Nunc) and grown at 37° for 5 h. The plate was washed with water using a microplate washer (Tecan Hydrospeed) and the Nluc working solution was diluted in PBS (30 µl PBS + 10 µl NLuc working solution) and added to each well to measure luminescence using the Tecan Spark reader with following settings: attenuation: auto, settle time: 0 ms, integration time: 100 ms.

For the inhibitor screen, 0.5 µl of each compound (Table S2) was added to 50 µl bacterial culture prior to incubation at 37°C for 5 h, and the plate was processed as described above.

### SDS PAGE, Western blotting and immunodetection

For protein detection, samples were separated by SDS PAGE using SERVAGel^TM^ TG Prime^TM^ 8-16% precast gels and transferred to a PVDF membrane (Bio-Rad) by standard protocols. Membranes were probed with primary antibodies α-SctP (39), α-SctE (39), α-c-Myc and α-SctJ (39). Secondary antibodies were goat anti-mouse IgG DyLight 800 conjugate. Detection was performed using the Odyssey imaging system (Li-Cor).

### MBP-NLuc and MBP-HiBiT expression and purification

NLuc and HiBiT, respectively, were cloned into a pMal-c5X vector to yield a translational fusion with maltose-binding protein (MBP). *E. coli* BL21 was transformed with the plasmids. Bacterial cultures were grown overnight at 37°C in LB broth and back-diluted in Terrific Broth (TB) the next day to an A_600_ = 0.1. They were grown to an A_600_ = 0.6-0.8 at 37°C. Subsequently, expression of MBP-NLuc/HiBiT was induced by addition of IPTG to a final concentration of 0.5 mM, after which bacteria were further grown at 37°C for 4 h. Bacterial cells were harvested by centrifugation (6,000 × *g*, 15 min, 4°C) and resuspended in column binding buffer (CB) containing 200 mM NaCl, 20 mM Tris-HCl (pH 7.4), 1 mM EDTA, Protease inhibitor (Sigma-Aldrich, P8849, 1:100), DNAse 10 µg/ml, 1 mM MgSO_4_ and lysozyme (10 µg/ml) and lysed with a French press. The obtained solution containing cell lysate and cell debris was two times centrifuged at 15,000 × *g* for 20 min at 4°C. MBP-NLuc/HiBit in the clear lysate was bound to an amylose resin (NEB), washed with CB and eluted by 10 mM maltose in the same buffer. Buffer was exchanged to PBS by using the Amicon Ultra system (Merck).

### Stability test of NLuc

40 µl Purified MBP-NLuc was added (2 µg, final concentration) to 1 ml LB/ 0.3 M NaCl and to 1 ml culture supernatant of wild type *S*. Typhimurium. Samples were kept either at 37°C, at room temperature, or on ice for up to 4 h. Aliquots were removed over time and transferred to a 384-well plate, 25 µl of the NLuc working solution was added and luminescence was directly measured in a microplate reader (Tecan Spark).

### Kinetic measurement

SipA-NLuc was introduced into the chromosome of *S*. Typhimurium, *P*ara-hilA by allelic exchange. The resulting strain was grown overnight at 37°C in LB/0.3 M NaCl, and was back-diluted the following day to an A_600_ = 0.1. Bacterial cultures grew to an A_600_ = 0.9 in an Erlenmeyer flask in a 37°C water bath, stirred with a magnet stirrer. Expression of *S*PI-1 was induced by addition of arabinose to a final concentration of 0.02% (v/v) and samples were taken at different time points thereafter for assessment of the luminescence of secreted SipA-NLuc or for immunodetection of SctJ. For testing the role of PMF inhibitors, bacterial cells were washed twice after reaching an A_600_ = 0.9 in LB/0.3 M NaCl containing either 120 mM Tris-HCl, pH 7.0 for CCCP (Sigma) or 120 mM Tris-HCl, pH 7.0 and 150 mM KCl for valinomycin (Sigma). For potassium benzoate, cells at the same growth stage (A_600_ = 0.9) were harvested and then washed twice with LB/0.3 M NaCl containing 80 mM MES buffer, pH 6.8. The cultures in the different media (without inhibitor, with 0.02% (v/v) arabinose) were kept in the water bath at 37°C and 200 µl of samples were taken at different time points and kept on ice. The inhibitors were added to the bacterial culture 60 min after *hilA*-induction. Cultures were kept in the water bath and samples were taken every 10 min. Samples were centrifuged to separate whole cells and supernatant. 25 µl of the supernatant was transferred to a white 384-well plate and luminescence was measured upon addition of the Nluc working solution in a luminometer.

### Generation of stable HeLa cell line expressing LgBiT

LgBiT was cloned into the MCS of pLVX-EF1α-IRES-Puro (Takara) resulting in pLVX-EF1α-LgBiT-IRES-Puro by Gibson assembly. 24 h prior to transfection, three 10 cm cell culture plates containing each 4 × 10^6^ HEK 293T cells in 8 ml DMEM supplemented with 10% FCS (v/v) and sodium pyruvate were incubated at 37°C, 5% CO2 overnight. The next day, 7 µg DNA of pLVX-EF1α-LgBiT-IRES-Puro in 600 µl sterile water was added to Lenti-X Packaging Single Shot (Takara). The containing pellet was completely resuspended and the solution incubated for 10 min at room temperature to allow formation of nanoparticle complexes. Finally the DNA/nanoparticle solution was added dropwise to the HEK 293T cells. After 4 h of incubation at 37°C, 6 ml growth medium was added and cell supernatant was harvested after 48 h and sterile filtered. In total 42 ml supernatant were reduced to a total volume of 4.2 ml used Lentix-Concentrator (Takara) exactly according to the protocol of the manufacturer. The viral suspension was aliquoted and stored at −80°C. The virus titer was determined using the QuickTiter Lentivirus Titer Kit (Cell Biolabs) according to the manufacturers protocol. The viral supernatant was then diluted to a final MOI of 2-10 in 10% FCS-VLE RPMI, supplemented with polybrene (4 µg/ml final concentration) and added to HeLa cells (5 × 10^5^ cell in 500 µl medium in six well plates). After overnight culture, medium was exchanged and cells were cultured for another day. The cells were then split, transferred to cell culture plates, and 2 µg/ml puromycin was supplemented. After outgrowth of stably transduced cells, single cell clones were generated by single cell dilution. Various cell clones were tested and verified for LgBiT expression by lysing the cells and performing a luciferase assay by the addition of purified MBP-HiBiT in the Hibit Lytic Buffer from the Hibit Lytic Detection Kit. Buffer and substrate was added in 1:50 ratio as described in the manufacturer’s protocol, MBP-Hibit (2 mg/ml) was added in 1:100 ratio to the buffer-substrate mixture.

### Injection assay and injection kinetics

1 × 10^4^ HeLa cells and HeLa LgBiT cells were seeded out in white 96 well plates with glass bottom 24 h before infection in 100 µl DMEM + 10% FCS (GIBCO). *S*. Typhimurium was washed and resuspended in HBSS to infect the cells at a MOI = 50 for 60 min. After infection, cells were gently washed with a microplate washer (Tecan Hydrospeed, 5 cycles dispensing and aspirating (speed: 70 µl/sec)) using 1 × PBS (GIBCO). A final wash volume of 100 µl was used together with 25 µl of Nanoglo live cell assay buffer (Promega) containing substrate for luminescence measurement in a Tecan Spark reader with the following settings: attenuation: auto, settle time: 0 ms, integration time: 1,000 ms. For monitoring the injection kinetics, HeLa LgBit cells were seeded out and *S*. Typhimurium bacteria in HBSS were added to the cells as described above. Directly upon addition of the bacteria, 25 µl of the reconstituted Nanoglo live cell buffer was added to the infection culture and luminescence reading was carried out for 90 min with a 2 min reading interval in the Tecan Spark with the same settings as for the injection assay.

## Acknowledgements

We thank Thomas Hesterkamp and Mark Brönstrup for continued input in high throughput assay development. We acknowledge receipt of the LgBiT/HiBiT split luciferase system by Promega before commercial release. This work was funded in part by the German Center for Infection Research (DZIF), grant TTU06.801 WP1.

## Supplemental Material

**Fig S1.**
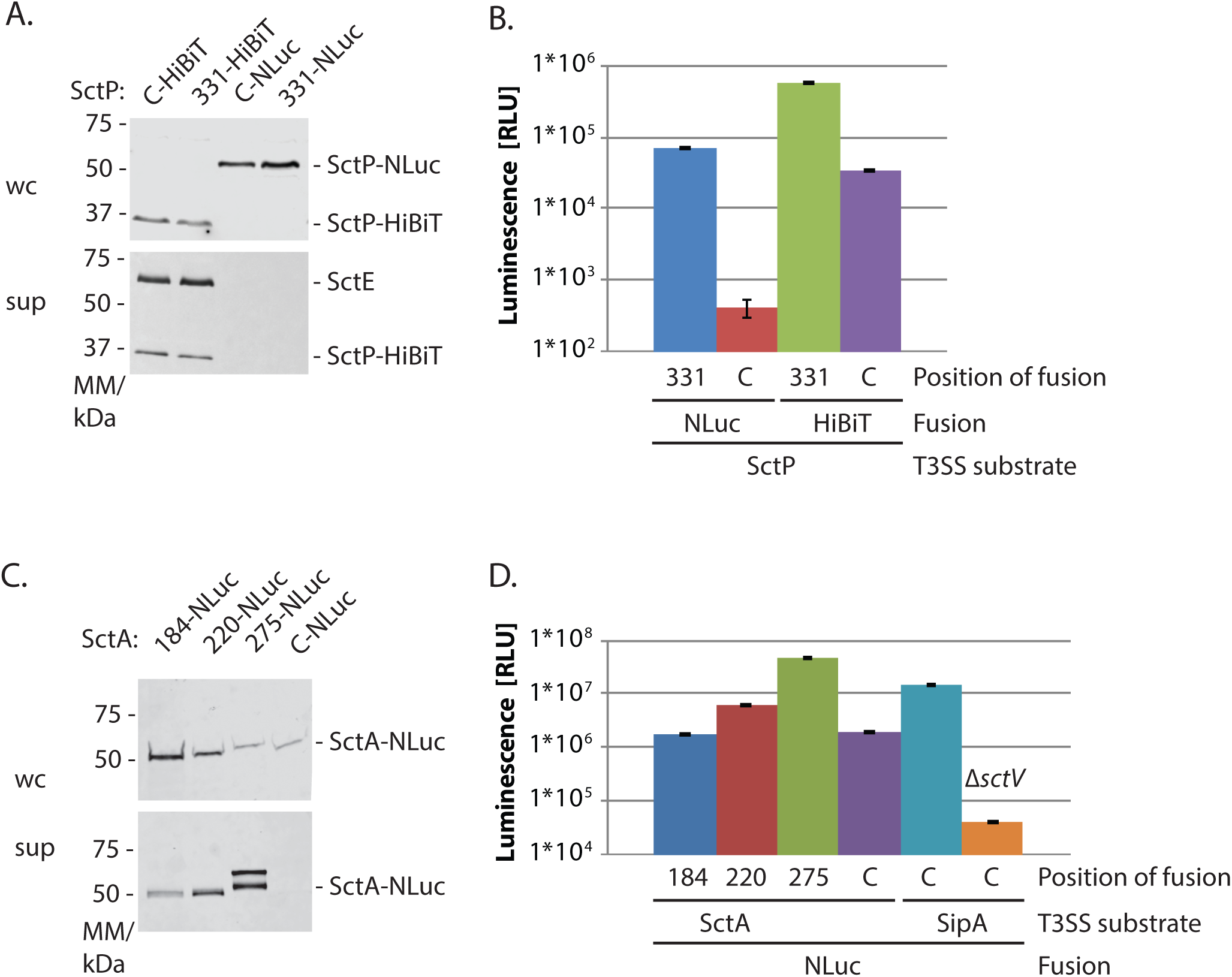
Expression and secretion of SctP-NLuc, SctP-HiBiT, and SctA-NLuc fusions. (A) Immunodetection of the indicated SctP-NLuc and SctP-HiBiT fusions, and of SctE on Western blot of SDS PAGE-separated culture supernatants and whole cell lysates. 331 means that NLuc or HiBiT was inserted behind residue 331 of SctP, so that the Shine-Dalgarno sequence of *sctQ*, which is overlapping with the gene of SctP, was unaffected. Note that SctE is not secreted when expressing SctP-NLuc fusions, i.e. SctP-NLuc cannot complement the needle length regulating function of SctP, thus substrate specificity switching to the secretion of intermediate substrates is not induced. (B) Luminescence of the indicated SctP-NLuc/HiBiT-fusions secreted into the culture supernatant. Data represent the mean (± standard deviation) of three technical replicates. Note that SctP_331_-NLuc can be detected in the culture supernatant by luminometry but not by Western blotting. Also note that split-NLuc generally gives lower luminescence than regular NLuc. (C) Immunodetection of the indicated SctA-NLuc fusions on Western blot of SDS PAGE-separated culture supernatants and whole cell lysates. The numbers (184, 220, 275) mean that NLuc was inserted behind these residues of SctA. The insertion positions where chosen based on the structure of *S*. Typhimurium SctA-1. Secreted SctA_275_-NLuc reproducibly appeared as a double band for unknown reasons. (D) Luminescence of the indicated SctA-NLuc and SipA-NLuc-fusions secreted into the culture supernatant. Data represent the mean (± standard deviation) of three technical replicates. Note that SctA_C_-NLuc can be detected in the culture supernatant by luminometry but not by Western blotting. Also note that internal fusions of NLuc are acommodated well, with SctA_275_-NLuc providing even stronger signal than SipA-NLuc. Abbreviations: sup: culture supernatant, wc: whole cell lysates, C: C-terminus, RLU: relative luminescence units, NLuc: NanoLuc luciferase, T3SS: type III secretion system

**Fig S2.**
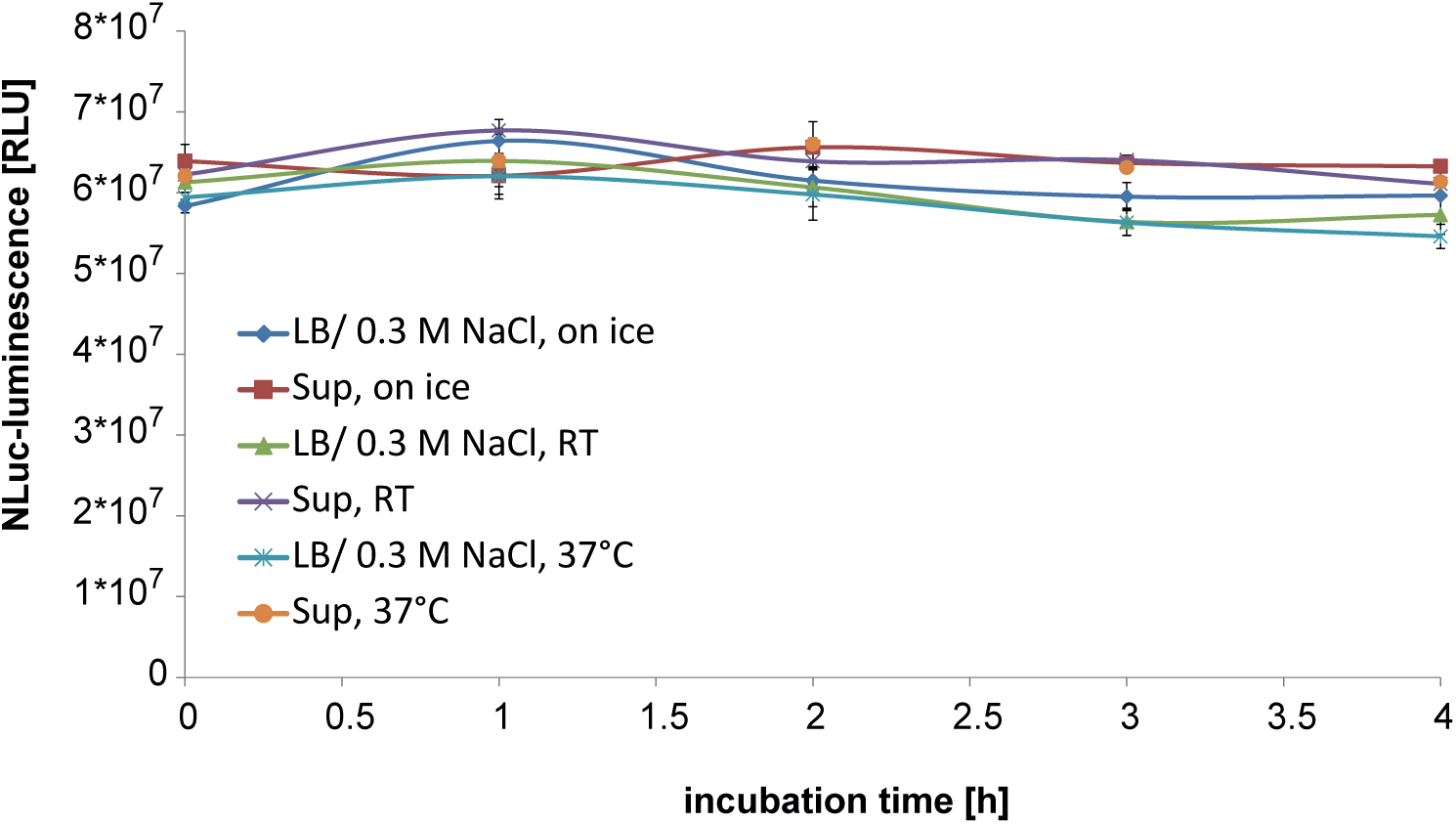
Stability of NLuc in LB/ 0.3 M NaCl and in culture supernatant. The enzymatic activity of purified NLuc was determined after incubation for 4 h at different conditions (on ice, room temperature (RT) and 37°C) in fresh LB/ 0.3 M NaCl and in filtered culture supernatant.

Table S1 Statistics of the reproducibility assessment of the 384-well microplate format NLuc secretion assay

Table S2 Layout of compound screening test plate incl. SipA-NLuc secretion of one measurement

Table S3 Strains, plasmids, primers

